# Intensity coded octopaminergic modulation of aversive crawling behavior in *Drosophila melanogaster* larvae

**DOI:** 10.1101/2020.09.04.281022

**Authors:** Florian Bilz, Madeleine-Marie Gilles, Adriana Schatton, Hans-Joachim Pflüger, Marco Schubert

**Affiliations:** Department of Biology, Chemistry and Pharmacy, Institute of Biology - Neurobiology, Freie Universität Berlin, Königin-Luise-Strasse 1-3, 14195 Berlin, Germany; Department of Biology, Chemistry and Pharmacy, Institute of Biology - Animal Behavior, Freie Universität Berlin, Takustrasse 6, 14195 Berlin, Germany

**Keywords:** functional calcium imaging, *in vivo*, ventral nerve cord, VUM neurons, noxious and gentle tactile stimulation

## Abstract

Activation and modulation of sensory-guided behaviors by biogenic amines assure appropriate adaptations to changes in an insect’s environment. Given its genetic tool kit *Drosophila melanogaster* represents an excellent model organism to study larger networks of neurons by optophysiological methods. Here, we studied stationary crawling movements of 3^rd^ instar larvae and revealed how the octopaminergic VUM neuron system reacts during crawling behavior and tactile stimulations. We conducted calcium imaging experiments on dissections of the isolated nervous system (missing all sensory input) and found spontaneous rhythmic wave pattern of neuronal activity in VUM neuron clusters over the range of thoracic and abdominal neuromeres in the VNC. In contrast, *in vivo* preparations (semi-intact animals, receiving sensory input) did not reveal such spontaneous rhythmic pattern. However, tactile stimulations activated different clusters of the VUM neuron system simultaneously in these preparations. The activation intensity of VUM neurons in the VNC was correlated with the location and degree of body wall stimulation. While VUM neuron cluster near the respective location of body wall stimulation were less activated more distant cluster showed stronger activation. Repeated gentle touch stimulations led to decreased response intensities, repeated harsh stimulations resulted in increasing intensities over trials. Optophysiological signals correlated highly with crawling behavior in freely moving larvae stimulated similarly. We conclude that the octopaminergic system is strongly coupled to the neuronal pattern generator of crawling movements and that it is simultaneously activated by physical stimulation, rather intensity than sequential coded. We hope that our work raises the interest in whole biogenic network activity and shows that octopamine release does not only underlie “the more the better” principle but instead has a more complex function in control and modulation of insect’s locomotion.

## Introduction

Insects use information about their environment and internal physiological states to execute fast, reflex-like sensory-guided behavior. In contrast to goal-directed behavior which depends on multimodal integration and memory in higher brain centers of the supraoesophageal (cerebral) ganglion (hereafter referred to as ‘brain’) some autonomous rhythmic behaviors such as crawling, walking, swimming or flying are generated brain-independently in the nerve cord and the gnathal ganglia. Here, neuronal networks, including motoneurons that innervate the muscles as output elements, are responsible for the execution of coordinated behaviors such as locomotion. While in the intact animal, central pattern generators (CPG) are modulated by sensory input and the regulatory control of the brain, in isolated ventral cords they can work autonomously. The importance and function of neuronal CPGs for locomotor behavior in *Drosophila* larvae is discussed extensively (Fox *et al.*, 2006; Song *et al.*, 2007; Xiang *et al.*, 2010; Berni *et al.*, 2012; Gomez-Marin & Louis, 2012). The modulation of CPGs by biogenic amines adds another important variable to the subject. In parallel to the motor network a small-sized network of octopaminergic neurons release octopamine (OA) onto neuromuscular junctions (NMJs) and potentially onto synapses within CPGs to modulate synaptic transmission and other physiological processes important for motor behaviors. In large insects such as locusts, the octopaminergic neurons of a thoracic ganglion, for example, are divided into different subpopulations with different morphologies (Watson, 1984; Duch *et al.*, 1999; Kononenko & Pflüger, 2007) and functions (Burrows & Pflüger, 1995; Baudoux *et al.*, 1998; Duch & Pflüger, 1999). Techniques to alter the concentrations of biogenic amines were used to exploit effects on individual or small groups of neurons and the behavioral consequences. However, how whole neuronal aminergic networks in insects are functioning system-wide is less investigated and difficult to achieve with electrophysiological recording methods but, as demonstrated here, possible by optophysiological methods.

Octopamine is a member of the phenolamine family, synthesized from the amino acid L-tyrosine functioning as a neurotransmitter, a neuromodulator and a neurohormone in invertebrates. Based on structural and functional similarities, the monoamine OA and its direct precursor tyramine (TA) constitute the invertebrate counterpart of the vertebrate’s norepinephrine and dopamine system (Roeder, 2005; Vömel & Wegener, 2008; Selcho *et al.*, 2012). Octopamine was first found and described in the salivary glands of octopoda (Erspamer, 1948) and was later comprehensively studied in arthropods and insects where it plays an important role in a variety of behavioral contexts (Roeder *et al.*, 2003; Roeder, 2005; Farooqui, 2012). It acts in the central as well as in the peripheral nervous system where it is released mainly under stress conditions, e.g. starvation (Wicher, 2007; Suo *et al.*, 2009), aggression (Baier *et al.*, 2002; Stevenson *et al.*, 2005) or foraging (Fussnecker *et al.*, 2006; Scheiner *et al.*, 2006). Among its versatile functions, OA can alter the efficacy of neuromuscular transmission (Walther & Zittlau, 1998), changing energy metabolism (Mentel *et al.*, 2003), having hyperglycaemic action (Candy, 1978; Zeng *et al.*, 1996) and inhibiting myogenic oviduct rhythms, accessory hearts and gut movements (Orchard & Lange, 1986; Papaefthmiou & Theophilidis, 2001). In addition, it is discussed as the “reward transmitter” of insects in associative learning paradigms (Hammer, 1993; Hammer & Menzel, 1998; Schroll *et al.*, 2006; Waddell, 2013). Importantly, OA was found to be involved in the activity and modulation of central networks for locomotion (Hoyle & Dagan, 1978; Sombati & Hoyle, 1984; Duch & Pflüger, 1999; Fox *et al.*, 2006; Pflüger & Duch, 2011). Its action had been studied extensively in large hemimetabolous insects such as locusts (for review see Bräunig and Pflüger (2001) but a number of studies were also concerned with the holometabolous *Drosophila* fly (Hoyer *et al.*, 2008; Vömel & Wegener, 2008; Zhou *et al.*, 2008; Busch *et al.*, 2009; Busch & Tanimoto, 2010; Selcho *et al.*, 2014; Pauls *et al.*, 2018), and the role of both amines, OA and TA in larval crawling (Selcho *et al.*, 2012) or OA in flight (Brembs *et al.*, 2007).

Common in insects, octopamine is released by a special class of neurons with bilaterally symmetrical axons in the ventral nerve cord and gnathal ganglia. These neurons have dorsal or ventral unpaired median cell bodies which, therefore, were called DUM or VUM neurons, respectively (Plotnikova, 1969; Crossman *et al.*, 1971; Hoyle, 1974; 1975; Hoyle & Dagan, 1978; Arikawa *et al.*, 1984; Bräunig & Pflüger, 2001). In locusts, these neurons fall into subpopulations which are differentially recruited during various motor tasks (Burrows & Pflüger, 1995; Baudoux & Burrows, 1998; Baudoux *et al.*, 1998; Duch *et al.*, 1999; Duch & Pflüger, 1999). In stick insects they are recruited during walking behavior (Mentel *et al.*, 2008) and descending VUM-neurons from the gnathal ganglion are involved in sensory gating (Stolz *et al.*, 2019). However, in the fruit fly, *Drosophila melanogaster*, it is unknown whether octopaminergic VUM neurons are also functionally subdivided during motor behavior. *Drosophila’s* 3^rd^ instar larval developmental state offers an octopaminergic system easily accessible for optophysiological investigations when compared to adult animals. Cell bodies of octopaminergic neurons can be found in the brain lobes and in the ventral nerve cord (VNC). As the larval body, the VNC is segmentally organized displaying three suboesophageal (s1-s3), three thoracic (t1-t3) and eight abdominal neuromeres (a1-a8), plus a small terminal neuromere at the end (a9). Clusters of octopaminergic cell bodies can be found in each neuromere. Thoracic and abdominal clusters can contain paired (VPM; t1-t3, a1) and unpaired (VUM; t1-t3, a1-a9) ventromedial neurons. Only the terminal neuromere possesses a dorsomedial cluster (DUM) (Vömel & Wegener, 2008; Selcho *et al.*, 2012) (Fig. 1A). The primary neurites of thoracic and abdominal VUM neurons are projecting first dorsally and then bifurcate, projecting laterally and leaving the VNC via the peripheral nerve strands (Fig. 3A and Suppl. Fig. 2A). The axonal VUM neuron projections target all the animal’s body wall muscles and modulate NMJ synapses by OA release (Monastirioti *et al.*, 1995; Monastirioti, 1999; Koon *et al.*, 2011; Koon & Budnik, 2012). Therefore, in this study we attempt to monitor the combinatory activity of the entire thoracic and abdominal population of VUM neurons in the VNC during larval crawling behavior simultaneously by optophysiological methods.

**Figure 1.**
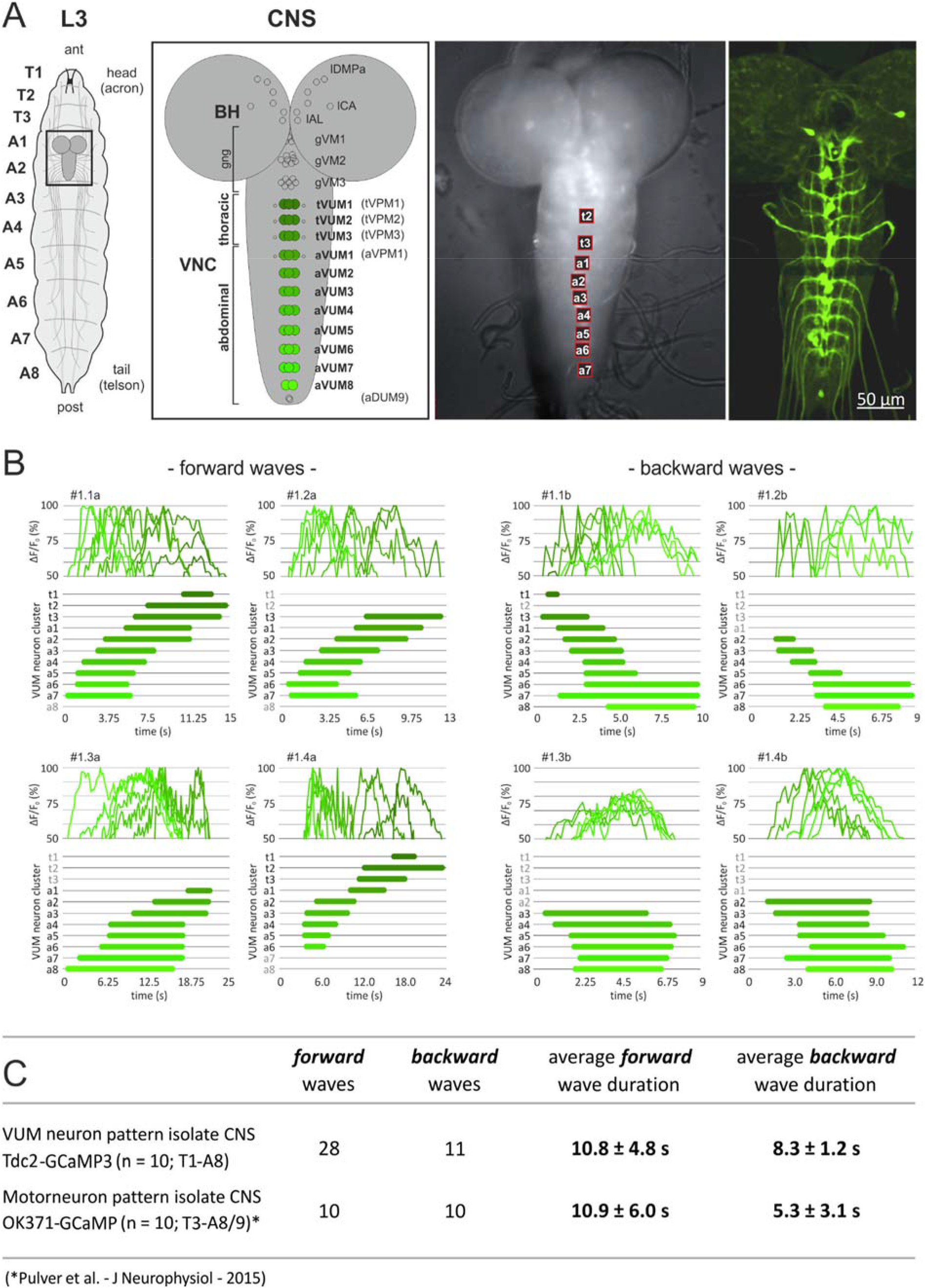
Spontaneously occurring activity wave pattern in octopaminergic VUM neuron cluster. Panel **A** shows the segmented body (T1-A8) of a 3^rd^ instar larvae and its CNS with neuronal projections innervating the body wall of each hemisegment (*left*). The CNS consists of two brain hemispheres and one elongated VNC. Three octopaninergic cell body cluster can be found in both brain hemispheres (lDMPa: larval anterior dorsomedial protocerebrum cluster; lCA: larval calyx cluster; lAL: larval antennal lobe cluster; after (Selcho *et al.*, 2014)) and 15 cluster in the VNC, three of which can be found in the gnathal (gVM1-3), three in the thoracic (tVUM1-3; tVPM1-3) and 9 in the abdominal neuromeres (aVUM1-8; aVPM1; aDUM9). Our investigation concentrated on the thoracic (tVUM1-3) and abdominal cluster (aVUM1-8) colored in different shades of green (*center left*). The anatomical grey scale image shows the ventral side of the brain. ROIs were set where cluster could be identified based on GCaMP5 auto-fluorescence (red squares, 5 × 5 pixel). In the shown example cluster tVUM2-aVUM7 (t2-a7) appeared in the focal plane and could be analyzed for fluorescence changes during imaging measurements (*center right*). The confocal microscopy image shows cluster and bilateral axonal projections of VUM neurons leaving each thoracic and abdominal VNC neuromere via paired nerve strands towards their appointed body segments (*right*). In panel **B** spontaneously occurring forward (*left*) and backward (*right*) wave pattern in the isolated brain preparations are visualized. Line graphs signify fluorescence changes over time passing the half maximum (50%) intensity threshold for each cluster. Beneath, horizontal bar graphs signify only the time fluorescence was passing the 50% threshold in each individual cluster to clearly visualize wave pattern. The table in panel **C** compares wave durations of forward and backward VUM neuron pattern with fictive locomotor pattern found in motoneurons of isolated brain preparations in *Drosophila* larvae by (Pulver *et al.*, 2015).

*Drosophila’s* 3^rd^ instar larvae offer a simple behavioral repertoire including peristalsis (forward and backward crawling), bending, turning and feeding (Green *et al.*, 1983; Fox *et al.*, 2006; Gomez-Marin & Louis, 2012). They use crawling behavior to execute escape reactions, to access food, to avoid competition and to find appropriate pupation areas. Forward crawling movements are initiated by the elevation and forward shift of the head and thorax body segments. Simultaneously a forward directed contraction wave starts traveling through the abdominal segments along the body axis from tail (body segment A8/9) to head (body segment A1) (Dixit *et al.*, 2008; Lahiri *et al.*, 2011; Berni *et al.*, 2012; Heckscher *et al.*, 2012). Backward movements are initiated by a retraction of the head and thorax followed by a backward directed contraction wave traveling through the abdominal body segments (Crisp *et al.*, 2008). All these behaviors are guided by sensory inputs, while the execution of movements (crawling and turning) are autonomous behavioral patterns created within the segmental motor-network of the nerve cord (Berni *et al.*, 2012). The larval body segments are targeted by these networks via nerve strands connecting the central nervous system (CNS = brain lobes + VNC) with the body periphery. These peripheral nerve strands carry not only motoneurons and efferent VUM neurons but importantly also the afferences from sensory receptor neurons. However, it is uncertain how exactly sensory input modulates the generation of rhythmic crawling behavior and, likewise, it remains an unclarified question, if, how and where the octopaminergic system might be involved in such modulation processes - only in the periphery or potentially also in the CNS?

In the present study we used tactile stimulations of different degrees in strength to initiate peristaltic crawling movements and to activate the animal’s octopaminergic system. By gentle and strong (noxious) physical touches against the external side of the animal’s body wall we aimed to activate different types of mechanoreceptors and studied the modulatory involvement of the octopaminergic system on the escape crawling behavior of the fruit fly’s 3rd instar larvae. We used calcium imaging techniques to examine the combinatory VUM neuron cluster activity in the VNC.

## Results

In different experimental approaches we investigated if and how octopaminergic neurons are involved in pattern generators responsible for larval crawling behavior in *Drosophila melanogaster*. Therefore, we used calcium imaging techniques to record neuronal activity signals from isolated (deafferented) CNS dissections in ***Experiment 1*** (n = 17) of 3^rd^ instar *Drosophila melanogaster* larvae expressing GCaMP3.0 in octopaminergic neurons (Tdc2-GAL4/UAS-GCaMP3.0). Furthermore, we used the same techniques and transgenic animals to record from semi intact *in vivo* preparations of the CNS in ***Experiment 2 and 3*** (n > 500). In these experiments ~60% of the isolated CNS dissections (n = 10) and in ~5% of the *in vivo* preparations (n = 27) we could measure changes in calcium concentrations signaling neuronal activity eligible for data analysis. In addition, we conducted behavioral open surface experiments observing the crawling performance of 3^rd^ instar larvae (n = 148) before and after tactile stimulations in ***Experiment 4***.

### Experiment 1. Spontaneous activity pattern in the VUM neuron system of isolated (deafferented) CNS preparations

We started our investigation by scrutinizing the role of octopaminergic neurons in active central pattern generators (CPG) of the isolated CNS. The isolation of the brain and ventral nerve cord prohibits any influence of afferent sensory input from the periphery on CPGs and permits only neuronal mechanisms and connections within the CNS. Besides arrhythmic neuronal activity, deafferented CNS dissections showed rhythmic wave pattern (posterior to anterior and *vice versa*) restricted to the thoracic and abdominal VUM neuron system in the VNC (Fig. 1B and Suppl. Movies 1 and 2). Rhythmic wave patterns are an indication for a possible participation of the octopaminergic system in larval crawling behavior since similar spatio-temporal patterns were already found in VNC motoneurons for a fictive crawling pattern (Pulver *et al.*, 2015). From 10 larval CNS dissections we analyzed 28 forward and 11 backward waves. Sixty percent of the animals showed both forward and backward waves whereas 40% showed only forward waves. Their occurrence was irregular and, in some cases, overlapping meaning that a new wave was initiated already before the preceding one was terminated. The average wave time lasted 9.8 sec ± 4.4 s.d. with forward waves being slightly slower (10.8 sec ± 4.8 s.d.) than backward waves (8.3 sec ± 1.2 s.d.) (unpaired t-test, p = 0.039) (Fig. 1C). The average VUM neuron activity duration for individual clusters was found 4.2 sec ± 2.5 s.d. In one exceptional case we were able to see activation pattern beyond thoracic and abdominal clusters in the gnathal ganglia cluster and the brain cluster. However, these activations could not be assigned to be part of any wave pattern. After recordings of spontaneous activity, we pharmacologically induced VUM neuron activity by applying pilocarpine (a muscarinic acetylcholine receptor agonist) to the CNS dissections. Although pilocarpine induces activity pattern in the CNS of many insects (Ryckebusch & Laurent, 1993; Büschges *et al.*, 1995; Johnston & Levine, 1996; Johnston *et al.*, 1999) its application neither changed the average wave time durations (10.3 sec ± 4.6 s.d.) nor increased the average cluster activity duration (4.7 sec ± 2.6 s.d.) (unpaired t-test, p > 0.05 in both cases). The maximum change in fluorescence intensity during wave pattern reached a value of 23.52 ± 9.3 (change in absolute grey scale values, calculated against the background).

### Experiment 2. Activity pattern in the VUM neuron system of in vivo preparations

In contrast to *Experiment 1* we were not able to see spontaneous neuronal activity pattern in the VUM neuron system of *in vivo* preparations. *In vivo* preparations of larvae had the brain and all nerve cords with afferent and efferent neurons completely intact. These larvae were able to execute stationary crawling movements in forward and backward direction while being held in place by pins at the anterior head segment and the telson (Fig. 2A and Suppl. Movie 3). These stationary movements occurred spontaneously without any experimental stimulations. Measured over 5 minutes, crawling or other behavioral movements were not accompanied by related neuronal activity wave pattern in the VUM neuron system as found in deafferented CNSs of *Experiment 1* (Fig. 2B, *left panel*). Thus, sensory feedback from stationary crawling movements is not sufficient to trigger VUM neuron activity or wave pattern in *Drosophila* larvae.

**Figure 2.**
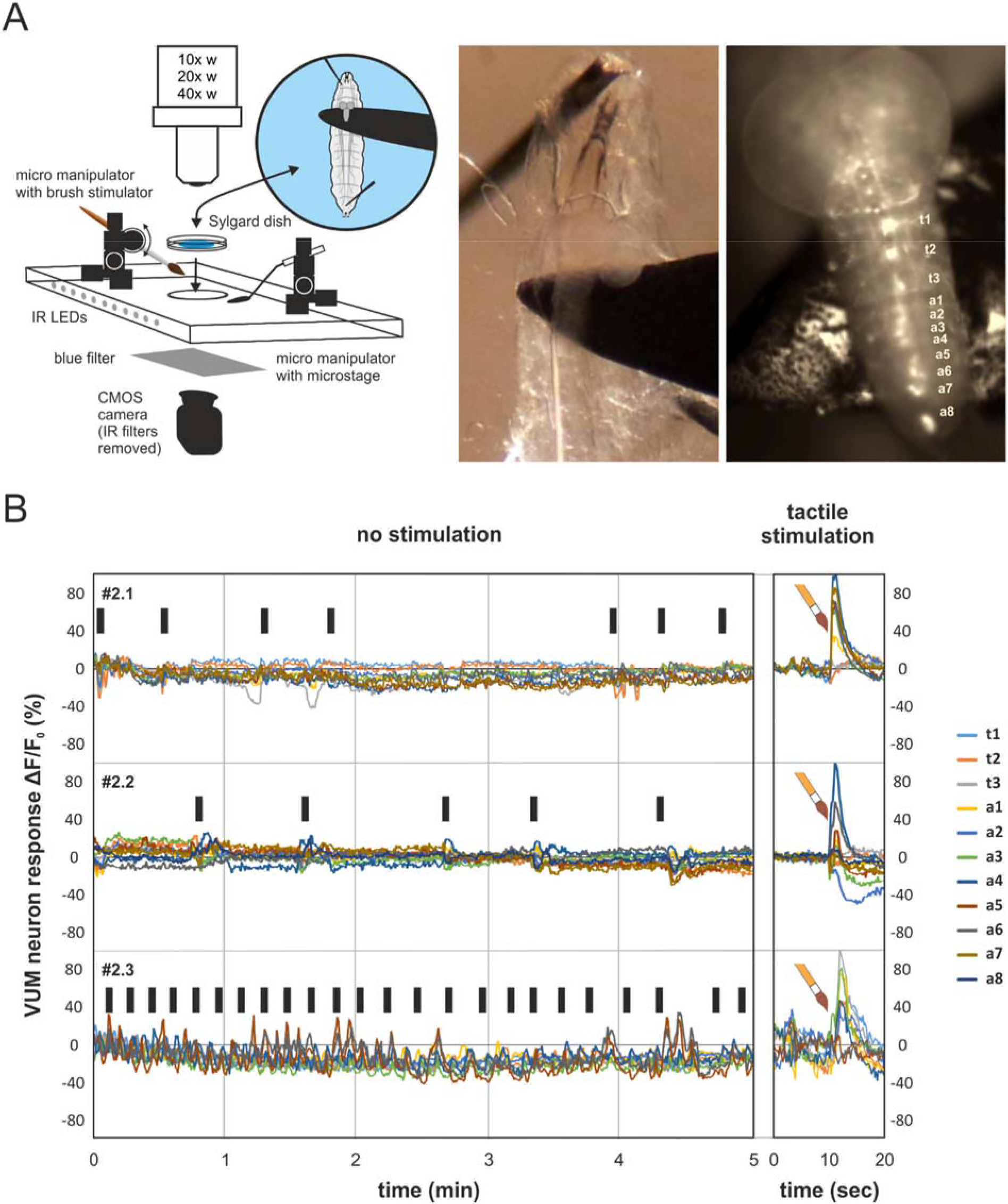
Activity pattern in the VUM neuron system of in vivo preparations. The sketch in panel **A** shows important parts of the experimental setup (*left*). Third instar larva were pinned onto a Sylgard dish and were covered by saline. The dish was placed under the microscope on a acryl glass platform equipped with two micromanipulators for stimulators and the microstage. The transparent platform had infrared (IR) LEDs and a CMOS camera allowing for video capturing of larval movements in the dark. The microstage was carefully positioned under the VNC (*center*) and the fluorescence of identified VUM neuron cluster were imaged (*right*). Panel **B** shows calcium imaging recordings over 5 minutes of VUM neuron cluster in the VNC of unstimulated larvae (*left*). Black bars signify larval crawling behavior confirmed by control ROIs outside the VNC in calcium imaging analysis and video capturing. Unstimulated larvae showed no consistent VUM neuron responses correlated to crawling movements. Sixty seconds after each 5 minute recording a new recording started comprising a tactile brush stimulation after 10 seconds (*right*). Tactile brush stimulations resulted in strong VUM neuron responses in many abdominal and thoracic cluster.

Conversely, when the body wall of a larva was stimulated by harsh tactile brush stimulation at the posterior end of the larvae the thoracic and abdominal VUM neuron system responded strongly. However, stimulations did not initiate wave pattern activity but led to strong simultaneous activity (in the boundaries of our time resolution given by the calcium sensor GCaMP3, t^1/2^ = 83 ± 2 ms) (Fig.2, *right panel*). These results were found consistently between animals. The maximum change in fluorescence intensity after harsh brush stimulation reached on average a value of 143.91 ± 28.27 (change in absolute grey scale values, calculated against the background).

In very rare cases pilocarpine application to intact brains (incubation of 5-10 min) led to flicker (on-off) activity in individual VUM neuron cell bodies lacking any coordination within the same or between different VUM neuron cluster (Suppl. Movies 4 and 5). Thus, individual VUM neurons have the potential to show independent activity from VUM neurons of the same or other cluster. However, the phenomenon of individual VUM neuron activity appeared only after pharmaceutical application or after damaging neuronal tissue during dissections.

### Experiment 3. VUM neuron pattern in response to consecutive tactile stimulations

To reveal how stable and consistent simultaneous VUM neuron responses occur after tactile stimulation we applied multiple stimulations within each experimental run. In addition to harsh brush stimulations (Fig. 3, *in shades of red* and Suppl. Movie 6) we also applied gentle rod stimulations (Fig. 3, *in shades of blue* and Suppl. Movie 7) to see if and how the VUM neuron system responds to different degrees of tactile stimulation and body wall deflections. Larvae were stimulated three times in each experimental run after 10, 30 and 50 seconds (20 seconds inter trial interval, ITI) either at posterior or anterior body segments.

**Figure 3.**
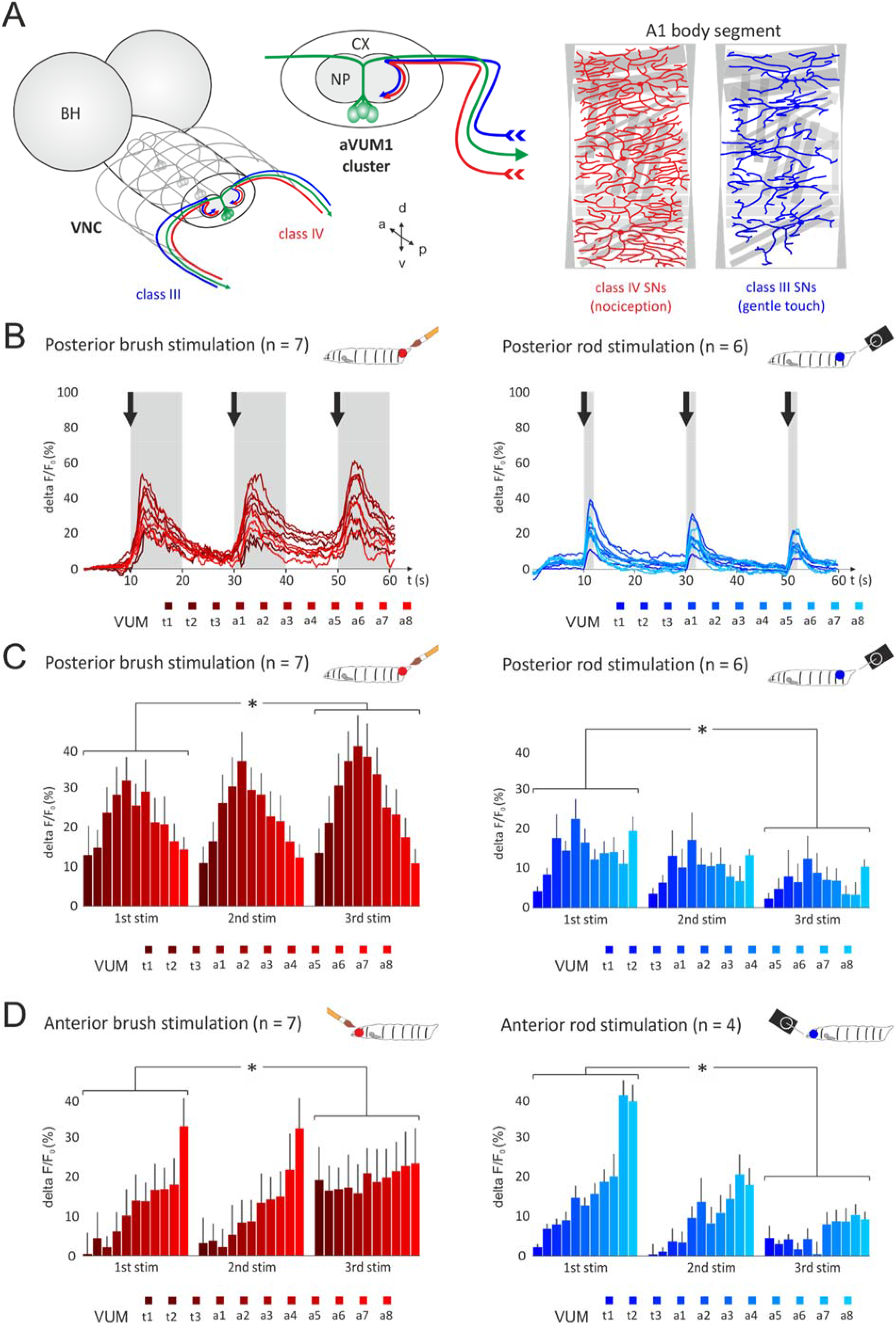
VUM neuron pattern in response to consecutive tactile stimulations. **A** Schematic drawing of the paired brain hemispheres and the VNC cut in the axial plane at abdominal VUM neuron cluster a1 (aVUM1). Three thoracic cluster (tVUM1-3) are shown in grey whereas abdominal cluster a1 is shown in green (*left*). VUM neuron cell bodies are located ventral medial and their primary neurite bifurcates dorsally descending axonal projections to corresponding body segments and NMJs. Ascending class III and class IV sensory neurons are depicted in blue and red, respectively. Class III and class IV neurons entre the neuropile at the dorsal part and project to their ventral target regions (*center*). Covering each body hemisegment harsh and gentle touch stimulations are perceived by class IV and class III sensory neurons, respectively (*right*). **B** Harsh (*left, in shades of red*) and gentle (*right, in shades of blue*) tactile stimulation of the larval body wall elicit different response activities in VUM neurons. The graphs are showing activation of identified thoracic and abdominal VUM neuron cluster over 60 seconds. Larvae were stimulated at the posterior end of the body either harshly with a brush (*left*) or gently with a small metal rod (*right*) three times in each experimental run after 10, 30 and 50 seconds (black arrows). Brush stimulations elicited strong and slightly increasing fluorescence responses over successive stimulations without total recovery of baseline fluorescence within 20 seconds after stimulations. Rod stimulations in contrast elicited strong but decreasing responses over successive stimulations. **C** Quantification of individual cluster responses to posterior brush stimulation (*left*) and posterior rod stimulation (right). After each stimulation responses to brush or rod stimulations were averaged over 50 (10 seconds) or 10 (2 seconds) consecutively imaged frames, respectively (as indicated by the grey boxes in B). A statistically significant difference between responses (over all 11 cluster) was found after the first and the third stimulation. **D** Quantification of individual cluster responses to anterior brush stimulation (*left*) and anterior rod stimulation (right) A statistically significant difference between responses (over all 11 cluster) was found after the first and the third stimulation. Error bars represent standard errors.

Posterior brush stimulations elicited strong and slightly increasing fluorescence responses over successive stimulations without total recovery of baseline fluorescence within 20 seconds ITIs (Fig. 3B, *left panel* and Suppl. Movie 8). For data analysis responses were averaged over 10 seconds after each stimulation and for each VUM neuron cluster. Although simultaneously active individual cluster differed in response strength, cluster aVUM8 (a8) - the cluster which gets sensory input from body segment A8 where the animal was stimulated the most - showed the lowest response intensity of all abdominal clusters. Cluster aVUM2 (a2) showed highest response intensity and all other abdominal clusters in between showed increasing intensities. However, after the peak response in cluster a2 response intensities decreased again in cluster a1, t3, t2 and t1. This pattern was consistently found in all consecutive stimulations. The average response intensity over all clusters were statistically compared between first, second and third stimulation (1^st^ stim, 2^nd^ stim and 3^rd^ stim, respectively). Paired t-tests found a significant increase between responses after the 1^st^ and 3^rd^ stimulation (p = 0.017; n = 7) (Fig. 3C, *left panel*).

Anterior brush stimulations elicited also strong and slightly increasing fluorescence responses over successive stimulations, however the simultaneous VUM neuron response pattern differed from posterior stimulations. Anterior brush stimulations led to weak responses in thoracic cluster and produced increasing response strength from abdominal cluster aVUM1 (a1) to a maximum in abdominal cluster aVUM8 (a8). Statistical comparison between responses after 1^st^, 2^nd^ and 3^rd^ stimulation using paired t-tests showed a significant increase between responses after the 1^st^ and 3^rd^ stimulation (p = 0.010; n = 7) (Fig. 3D, *left panel*). The maximum change in fluorescence intensity after tactile brush stimulation (including posterior and anterior stimulations) reached an average value of 104.48 ± 50.79 (change in absolute grey scale values, calculated against the background).

Posterior rod stimulations in contrast, elicited strong but decreasing responses over successive stimulations (Fig. 3B, *right panel* and Suppl. Movie 9). Responses were averaged over 2 seconds after each stimulation and for each VUM neuron cluster. Again, VUM neurons responded simultaneously and the response strength differed between individual cluster. Cluster aVUM8 (a7) - the cluster which gets sensory input from body segment A7 where these animals were stimulated - showed the lowest response intensity of abdominal cluster. Cluster aVUM2 (a2) showed highest response intensity and intensities decreased again in cluster a1, t3, t2 and t1. Paired t-tests found a significant decrease between responses after the 1^st^ and 3^rd^ stimulation (p < 0.001; n = 6) (Fig. 3C, *right panel*).

As for brush stimulations anterior rod stimulations produced weakest responses in anterior cluster and strongest responses in posterior cluster but showed decreasing responses over successive stimulations. Paired t-tests found a significant decrease between responses after the 1^st^ and 3^rd^ stimulation (p = 0.011; n = 4) (Fig. 3D, *right panel*). The maximum change in fluorescence intensity after tactile rod stimulation (including posterior and anterior stimulations) reached an average value of 52.23 ± 40.61 (change in absolute grey scale values, calculated against the background).

Altogether, VUM neurons showed consistently weaker responses in those clusters which are getting sensory input from stimulated body segments and stronger responses in clusters getting input from segments further away from stimulated segments.

### Experiment 4. Tactile stimulation of freely moving larvae - a correlation between optophysiological recordings and behavioral observation data

In a last step we wanted to investigate if tactile stimulations and hence a change in VUM neuron activity (as shown in *Exp. 2* and *Exp. 3*) have any effect on the larval crawling behavior. Since VUM neuron activity results in a release of octopamine at the target areas of axonal projections, the NMJs and muscle fibers (Fig. 4A), synaptic properties and muscle conditions are expected to be altered leading to changes in crawling dynamics (compare Koon et al. 2011).

**Figure 4.**
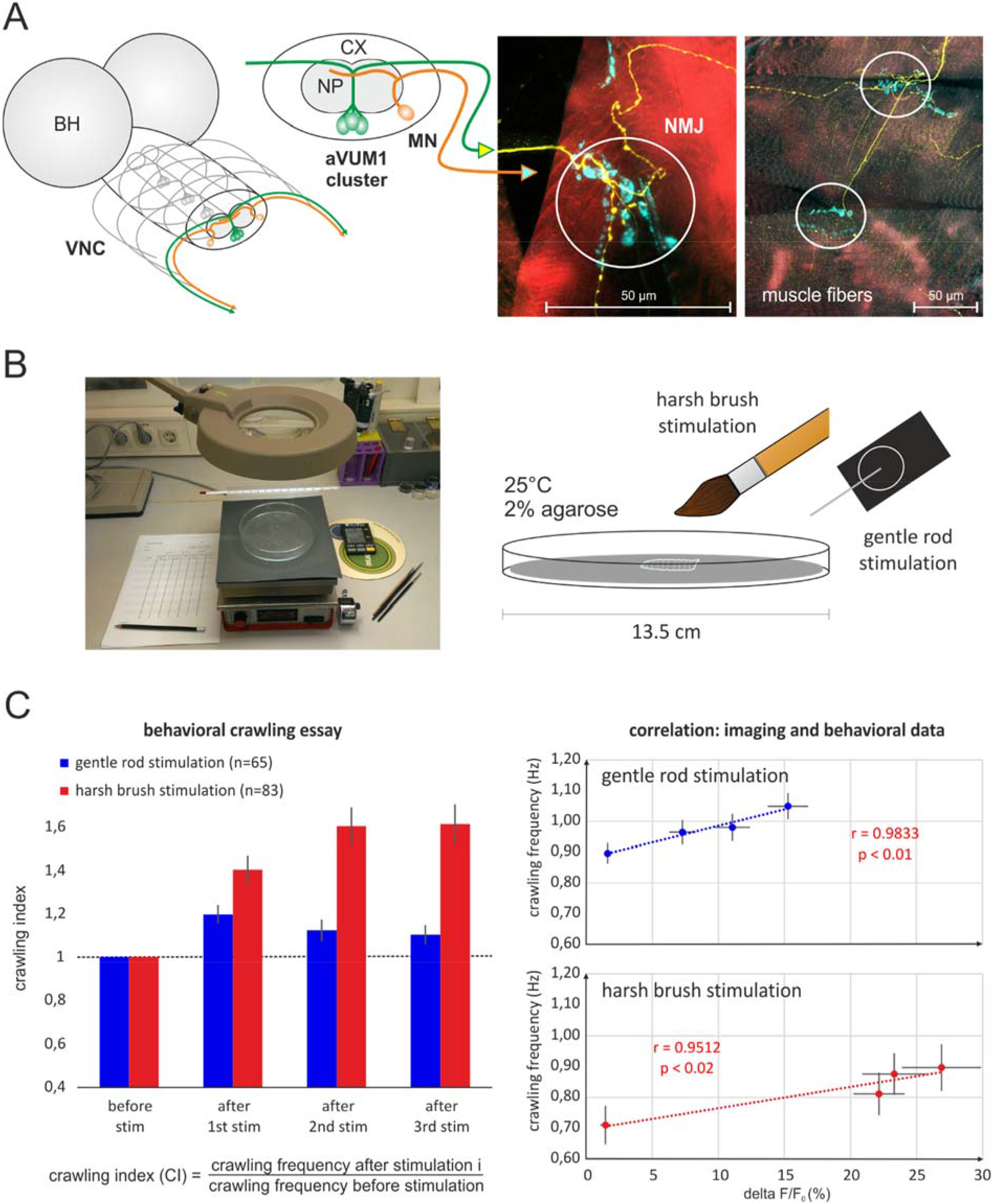
Tactile stimulation of freely behaving larvae. **A** Schematic drawing of the paired brain hemispheres and the VNC cut in the axial plane at abdominal VUM neuron cluster a1 (aVUM1) (left). Descending axonal projections of aVUM1 neurons (green) and motoneurons (orange) leave the VNC in parallel via the same nerve strands bilaterally and project to muscle fibers of their corresponding body segment A1. VUM neurons target NMJs at multiple muscle fibers and have lateral Type II projections along the muscle fibers (*right, confocal images*). In confocal images VUM neurons are depicted in yellow, NMJs in cyan and muscle fibers (actin filaments) in red. **B** In behavioral essays larvae were released in the center of a glass dish cast with 2% agarose. The dish was placed on a temperature controlled plate and behavioral performances were observed under a magnification glass (2x). Crawling events were counted before and after harsh brush and gentle rod stimulations in respective experimental groups. **C** Bar graphs show the crawling performance index of larvae during a time period of 20 seconds before any stimulation and 20 seconds after each of the three successive stimulations (ITI of 20 seconds). In blue and red responses to gentle rod (n = 65) and harsh brush (n = 83) stimulations are given, respectively. Responses to gentle rod stimulations increased the crawling frequency after the first stimulation but then decreased again after the second and third stimulation. In contrast, responses after harsh brush stimulations show ever increasing crawling frequencies after each stimulation (*left*). Correlation analysis between imaging and behavioral data were performed using linear regression analysis and Pearson’s correlation coefficients. Both data sets are highly correlated (*right*). Error bars represent standard errors.

In very simplistic behavioral assays we concentrated on changes in the crawling frequencies of larvae after repeated tactile stimulations of different degrees (gentle rod and harsh brush stimulations). Both types of stimulations were conducted in respective experimental groups. Before stimulations larvae in the gentle rod stimulation group displayed undisturbed foraging behavior with a crawling frequency of 0.90 Hz during 20 seconds (n = 65). After three successive stimulations with an inter trial interval (ITI) of 20 seconds frequencies first increased to 1.05 Hz and then decreased again to 0.98 Hz and 0.96 Hz after the 1^st^, 2^nd^ and 3^rd^ stimulation, respectively (Suppl. Fig. 3A). Larvae in the harsh brush stimulation group initially showed a crawling frequency of 0.71 Hz before stimulations (n = 83). After successive stimulations frequencies increased to 0.81 Hz, 0.88 Hz and 0.90 Hz after the 1^st^, 2^nd^ and 3^rd^ stimulation, respectively (Suppl. Fig. 3B).

To compensate for the difference in initial crawling frequencies (before 1^st^ stimulation) between experimental groups a crawling index was calculated before statistical comparisons were made. For index calculations in both experimental groups, the number of crawling events after each stimulation were divided by the number of crawling events before the 1^st^ stimulation. Therefore, controls before stimulation have a crawling index of 1. After the 1^st^ stimulation the index of both experimental groups increased, however stronger in the harsh brush stimulation group (gentle rod: 1.20; harsh brush: 1.40). After the 2^nd^ stimulation the index for the gentle rod stimulation group dropped to 1.12 whereas the index for the harsh brush stimulation group further increased to 1.60. After the 3^rd^ and last stimulation the index for the gentle rod stimulation group further decreased to 1.10 and it further increased in the harsh brush stimulation group to 1.61 (Fig. 4C, *left panel*, gentle rod group in blue and harsh brush group in red).

For correlation analysis between behavioral crawling frequencies and optophysiological signal intensities (averaged over all measured VUM neuron cluster) in both experimental groups we used linear regression analysis and Pearson’s correlation coefficients. Analysis revealed a significant correlation between both data sets in both experimental groups (gentle rod: r = 0.98, p < 0.01; harsh brush: r = 0.95, p < 0.02) (Fig. 4C, *right panel*). Therefore, successive tactile stimulations of different degrees lead to characteristic adaptation of both the neuronal VUM neuron activity and the crawling behavior. In case of gentle rod stimulations this means habituation after an initial increase and sensitization in case of harsh brush stimulations.

## Discussion

Motivated by the many electrophysiological studies concentrating on individual or small populations of VUM neurons simultaneously, we wanted to reveal how the VUM neuron system reacts during crawling behavior as an entity. Therefore, we conducted calcium imaging experiments on the nervous system of *Drosophila* larvae. In isolated larval CNS dissections that do not receive sensory input we found spontaneous rhythmic wave pattern of neuronal activity in VUM neuron clusters over the range of thoracic and abdominal neuromeres in the VNC. Surprisingly, *in vivo* preparations that receive sensory input did not show such spontaneous rhythmic pattern, no matter if the animals were executing crawling movements or not. However, tactile stimulations activated different clusters of the VUM neuron system simultaneously while the activation intensity was correlated with the location and degree of body wall stimulation. While the response of VUM neurons in stimulated neuromeres was low, it increased in distant neuromeres. Moreover, the response intensities to repeated gentle touch stimulations decreased, while they increased with repeated harsh stimulations over trials. The optophysiological signals correlated highly with crawling behavior in freely moving larvae stimulated similarly with gentle and harsh touches.

VUM neuron patterns found in our isolated CNS preparations were very similar to fictive crawling patterns in motoneurons of deafferented larval CNS of *Drosophila melanogaster* (Pulver *et al.*, 2015). This suggests a possible coupling of VUM neurons to the central pattern generator for larval crawling behavior. However, in many cases the isolated CNS produces rhythms only similar and not identical to the ones of intact animals but if phase relations of antagonistic systems are retained, the respective motor behavior of the isolated CNS is called “fictive”. Thus, removing sensory organs could lead to changes in the functional principles of the whole pattern generator (CPG + sensory organs) (Bässler, 1986). How sensory information from the periphery is processed in the VNC and the brain to allow smooth locomotor behavior is still a matter of extensive research. Here we propose VUM neurons as one possible system to modulate either indirectly the endogenous input or directly the central pattern generating neurons as a response to external tactile stimulation of the body wall. Two scenarios can be considered: first, tactile stimulation *triggers* the pattern generator provoking a reflex and second, tactile stimulation *modulates* the frequency of the already oscillating pattern generator. Both scenarios can be explained by an endogenous input gradually changed and pushed above threshold by sensory and/or modulatory neurons (e.g. octopaminergic VUM neurons) without input from higher brain centers. In this model, octopaminergic neurons would be involved in gating processes similar to what has been found in stick insects (Stolz *et al.*, 2019) or, perhaps, in changes associated with an animal being active and starting a dynamic period of behavior.

### Sequential modulation in VUM neuron cluster activity

The repeated sequential activation patterns in isolated CNS could have been triggered by either a sequential recruitment by neurons presynaptic to VUM neurons and/or by a delayed coupling between clusters. Work on *Manduca sexta* larvae showed that fictive crawling motor patterns were accompanied by rhythmic activation of unpaired median (UM) neurons in all segmental ganglia derived partially from a source within the gnathal ganglion (Johnston & Levine, 1996; Johnston *et al.*, 1999). Because of very short anterior-to-posterior activation delays (~10 ms) only one or few descending neurons of the gnathal ganglion projecting posteriorly to all segmental ganglia could be responsible for all efferent UM neuron inputs. However, the sequential activation in isolated CNS preparations of *Manduca sexta* larvae appear much faster than those found in *Drosophila melanogaster* larvae preparations. The reason for this could be differences in the network organization and/or neuronal properties but is so far unknown. In strong contrast to our isolated CNS preparations, VUM neuron activation in our *in vivo* experiments (whole animal with intact sensory system) was synchronous and appeared only after tactile stimulation, not during crawling. A possible reason for that could be the fixed position of the larvae allowing only stationary crawling movements. Importantly, under fixed conditions VUM neuron activation appears rather intensity than sequentially modulated. (Hu *et al.*, 2017) identified a pair of interneurons (a08n) with their cell bodies located in abdominal neuromere a8 and having ascending ramifications over all thoracic and abdominal neuromeres. These ramifications were situated in very close vicinity to nociceptive receptor neuron axon terminals in the VNC which can detect harsh mechanical stimulations of the larval body wall (class IV dendritic arborization neurons, class IV da). Importantly, a08n neurons responded to an optogenetically activation of these nociceptive receptor neurons. Thus, in principle, a08n neurons could spread information of tactile stimulation over the VNC with minimal delay and activate almost simultaneously a downstream octopaminergic network system wide. Although still unclear, mechanisms of an intensity modulation by the VUM neuron system could lead, in theory, to an aminergic readout about where and how strongly in the periphery the sensory system was activated, thus encoding a somatotopic map of the larval body wall. If VUM neurons and motoneurons are integral parts of the same pattern generator for fictive crawling behavior, it is noteworthy that simultaneous activity in VUM neurons would lead to octopaminergic modulation at their target NMJs preceding motoneuronal input at most of the body wall segments and muscles. This is however not contradictive because of the long-lasting effect of octopamine, resulting e.g. in synapses being prepared beforehand the arrival of neuronal input. The idea of a synchronous activation by a common interneuron is also promoted by recordings after pilocarpine application in which we saw cluster and individual VUM neurons activated individually. We expect subpopulations of VUM neurons to be recruited for the modulation of asymmetric or other more complex rhythmic or singular muscle activation patterns (e.g. rolling, turning or hunch behaviors) as already found in other insects (Duch & Pflüger, 1999; Pflüger & Duch, 2011). Furthermore, it would be most interesting to compare larval VUM neuron activity patterns with patterns of octopaminergic neurons in the adult systems during e.g. walking and flight behavior.

### Intensity modulation in VUM neuron cluster activity

The absence of successive VUM neuron cluster activity in *in vivo* preparations (animals perceiving sensory feedback) promotes the idea that it is not essential for the execution of basic crawling movements but, however, for their modulation. Simultaneous and intensity modulated VUM neuron activity was induced by body wall stimulation. In the periphery, octopaminergic neurons innervate the major part of body wall muscles (Monastirioti *et al.*, 1995) where octopamine increases presynaptic vesicle release (Koon *et al.*, 2011) at neural muscular junctions (NMJs) and contractions directly of body wall muscle fibers/cells (Ormerod *et al.*, 2013). Importantly, the effects are cell selective and correlate with the distribution of octopamine receptors (El-Kholy *et al.*, 2015) on different muscles within each body wall segment. Hence, octopamine has an important role as neuromodulator and a great potential impact on the coordination of different transversal and longitudinal muscles important for crawling activity (Ormerod *et al.*, 2018). Fox *et al.* (2006) revealed that mutants (*T*β*h*^*nM18*^) with a reduced level of octopamine and an increased level of tyramine exhibit fewer rhythmic neuronal bursts in motoneurons associated with body wall contraction waves. The same mutant animals lack crawling speed and make more pauses compared to wild type (Saraswati *et al.*, 2004; Schützler *et al.*, 2019). Our findings show that activity within the abdominal and thoracic VUM neuron system in the CNS is synchronous and intensity modulated as, in consequence, should be the octopamine release at NMJs in the body wall at the periphery. Together these findings suggest that octopamine release is important 1) for the coordination of muscle activity within an individual body segment, and 2) for the coordination between all body segments since octopamine is released in different intensities depending on the segment’s location. In our study octopamine was only released after physical stimulation, an event which in the natural environment represent a potential threat to the animal and should be behaviorally responded to with avoiding behaviors and an increased crawling speed for body displacement. This hypothesis is confirmed by our open field crawling behavior experiments, in which contraction frequencies increased after physical stimulations. This is consistent with the idea that octopamine is released to meet the coordinatively more challenging needs of “high” speed crawling movements. Conclusions about central and peripheral observations both strengthen our idea of octopamine as integral part of the pattern generator and modulator which increases muscle coordination and movement performance over all body segments. Sujkowski *et al.* (2017) reported that octopaminergic neurons mediate exercise adaptations increasing endurance of e.g. flight and climbing performances in adult *Drosophila* flies. In this case adaptations were not limited only to skeletal muscles but also e.g. to cardiac stress resistance. Octopamine mutants are also more vulnerable to environmental stress (Chentsova *et al.*, 2002) while wildtype flies increase octopamine levels in response (Hirashima *et al.*, 2000) suggesting that the effects are not only local but also systemic adapting the whole system to higher performances. Future studies on the effects of octopamine release like the coordination of muscles and body wall segments in the periphery, as well as central effects and systemic adaptation are needed to fully evaluate the specific role of octopamine and other neuromodulators and neuropeptides in larval and adult behavior (Ormerod *et al.*, 2018).

### VUM neuron responses in multiple trial stimulations

The induction of multiple tactile stimulations of two different intensities changed VUM neuron response patterns differently over trials. Different intensities of tactile stimulation activated probably two different classes of sensory neurons (class III and IV da neurons) resulting in different VUM neuron behaviors. Multiple gentle rod stimulations provoked initially an increase in VUM neuron activity after the first trial which decreased and adapted quickly in consecutive stimulations. However, on a different time scale this provides a similar effect as found by (Yan *et al.*, 2013) for class III da neurons which showed a quick adaptation to prolonged gentle touch stimulation. Therefore, the adaptation effect found in VUM neuron responses is probably caused by sensory input from adapting class III da sensory neurons. To what degree a downstream network and the VUM neurons themselves are involved in adaptation effects is unclear and needs further investigation. Harsh brush stimulations provoked initially a comparable increase in VUM neuron activity, however, after repeated stimulations we found VUM neuron activity increased rather than decreased. Because of the long livability of increased calcium levels, we think that brush stimulations are perceived by class IV da sensory neurons sensitive to noxious stimuli. Long lasting calcium increase indicates a mechanism to down regulate the neuronal activity threshold and thereby an increase of excitability and sensitivity. Since octopamine release at the NMJs increases the efficacy of synaptic transmission, a repeated noxious stimulation is responded by a continual increase of octopamine release, and an ever facilitated transmission at the NMJs. Additionally, it appears that a repetitive noxious stimulation of constant intensity can result in an increasing response strength of nociceptive neurons (centrally mediated sensitization) leading to even stronger escape behaviors after a second or a third noxious stimulation (Walters *et al.*, 2001; Ji *et al.*, 2003; Hu *et al.*, 2017; Tabuena *et al.*, 2017). Such an enhancer effect could be realized not only by increased sensitivity of nociceptors but also within downstream interneurons or parallel VUM neurons. Our behavioral data strengthens these interpretations since the same stimulations used to activate the VUM neuron system in imaging recordings were used to stimulate freely crawling larvae. Responses to multiple trial gentle rod and harsh brush stimulations followed exactly the same response pattern as VUM neurons did. Initial stimulations increased crawling frequencies for both kinds of stimulations, however, after the second and third rod stimulation crawling frequencies decreased whereas frequencies after brush stimulations increased. The strong correlation of optophysiological and behavioral data is striking and promotes the idea of a direct link between them.

## Material & Methods

### Animals and dissection

We used transgenic 3^rd^ instar larvae of *Drosophila melanogaster* expressing the calcium detector GCaMP3.0 in Tdc2 positive neurons (homozygous Tdc2-GAL4/UAS-GCaMP3.0) allowing optophysiological activity surveillance of tyraminergic/octopaminergic neurons. GCaMP3 rise half-life time t^1/2^ = 83 ± 2 ms; decay half-life time t^1/2^ or tau τ = 610 ± 32 ms (Tian *et al.*, 2009). Flies were reared at 25°C and raised on standard cornmeal/molasses media, at a 12/12 h light/dark cycle. For isolated brain imaging experiments larvae were dissected in cooled saline (85 mM NaCl, 20 mM KCl, 4 mM MgCl2, 2 mM CaCl2, 10 mM HEPES, 20 mM Sucrose in Aqua dest., pH 7.25) using forceps to extract the CNS from the body. Briefly, we held the larvae in place by one pair of forceps gripping it halfway down its body length and extracting the CNS by pulling at the mouth hooks with a second pair of forceps. After cleaning the brain tissue, it was pinned onto a Sylgard dish (Fig. 1A). For *in vivo* imaging experiments 3^rd^ instar larvae were anesthetized on ice before being pinned onto a Sylgard dish using one pin at the anterior head segment and a second pin at the telson (Fig. 2). In that way body displacement was restricted to peristaltic muscle contractions typical for larval crawling behavior. Subsequently, larvae were dissected in cooled saline. For animal dissections a shallow dorsal incision between the main tracheae was made starting between body segment A3 and A4 and running along the dorsal midline longitudinally up to the head segment. The left and right body walls were pinned flat to the bottom of the Sylgard dish. The esophagus, proventriculus and parts of the gut and glands were removed. The brain lobes, VNC, and segmental nerves were left intact. The Sylgard dish was transferred to a custom-made acryl glass microscope stage equipped with two micromanipulators enabling the precise adjustment of tactile stimulators and the ‘microstage’ (Fig. 2A). The microstage, a small custom-made metal spoon, was put under the VNC and was slightly elevated to minimize artefacts during body wall stimulation or larval crawling movements. The surface of the spoon was lacquered matt black to enhance contrast to the whitish transparent color of the VNC. The transparent microscope stage was further equipped with infrared (IR) LEDs to allow for video recording of larval movements in darkness from beneath and through the Sylgard dish. For video capturing we used a CMOS camera (UCMOS 03100 KPA, ToupTek Photonics, Hangzhou, China) with its IR filter removed and covered by a glass filter blocking wavelengths used for imaging recordings (Fig. 2A).

### Stimulation

Mechanical stimulations of the larvae’s body wall were applied by either a paint brush (round, size 1) or a small metal rod (0.04 cm in diameter) at the anterior or at the posterior body segments. In imaging experiments the brush was fixed to a rotating servomotor (SM-S2309S) controlled by an Arduino UNO (microprocessor developer board) which was externally triggered by imaging software. The brush was positioned laterally to the animal’s body wall (not touching it). When triggered it was swinging towards the body wall delivering a harsh mechanical (noxious) touch deflecting the body wall more than 40 μm (Yan *et al.*, 2013). The custom-made rod stimulator was composed of a miniature loudspeaker (LSM-S20K, EKULIT), to which a blunt metal rod was adhered. The loudspeaker was connected to an electric pulse generator (SD9 Stimulator, Grass Products, Warwick, Rhode Island, U.S.A.). By generating rectangular impulses, the diaphragm of the loudspeaker was vibrating and hence the metal rod was displaced. Intensity and duration of the impulse were regulated by the stimulator settings. The metal rod was positioned laterally to the animal’s body wall touching only one body hemisegment. A gentle mechanical touch (deflection of < 40 μm) (Yan *et al.*, 2013) was provoked at 100 Volt with a duration of 100ms. Each stimulation consisted of 5 body wall touches at 5Hz. The pulse generator was externally triggered by imaging software.

For pharmacological stimulation of the VUM neuron system we applied 5 μl of Pilocarpine (10^−4^ molar; 0,0244mg/ml) onto the *in vivo* brain dissection. Pilocarpine, a muscarinic agonist which mimics the action of neurotransmitter acetylcholine by binding muscarinic acetylcholine receptors is not hydrolyzed by the acetylcholinesterase.

### Functional in vivo calcium (Ca^2+^) imaging

We used an upright fluorescence microscope, Axioskop FS (Zeiss, Germany), equipped with a 10x (UMPlanFL 10x/0.30 w), 20x (XLUMPlanFL 20x/0.95 w) and 40x (LUMPlanFl 40x/0.80 w) water immersion Olympus objective. A Polychrome II (xenon-lamp, 75 W, Photonics, Planegg/Germany), provided light excitation at 475 nm and a filter set (Omega Optical, Vermont, USA) ensured passage of only relevant wavelengths (dichromatic mirror: DCLP500, emission filter: LP515). The emitted light was captured by a 12 bit CCD camera (SensiCam QE, pco.imaging, Germany) with a symmetrical binning of 4 (1.25×1.25 μm/pixel). For imaging measurements, a series of either 300 or 600 images (henceforth frames; 344×260 pixel) was recorded at 5 or 2 Hz, respectively. Longtime and low frequency measurements were used for observations without stimulations whereas short time and high frequency measurements were used for multiple stimulation trials. When stimulated, three successive stimulations with an inter stimulus interval (ISI) of 20 seconds were applied at 10, 30 and 50 seconds into the recording (frame 50, 150 and 250, respectively). All imaging data were analyzed using custom written software in IDL (Interactive Data Language, ITT Visual Information Solutions). A bleaching correction was applied for each frame by subtracting the median fluorescence from each pixel. An automated movement correction compensated for movement artifacts occurring during image sequence recordings. For data acquisition, we set regions of interest (ROIs) for each identifiable VUM neuron cell body cluster in the ventral nerve cord (VNC) of each animal. Mean values of each ROI (5×5 pixels per cluster) were calculated in each image (frame) of the recorded sequence. A data matrix was generated for the fluorescence changes of each identified cluster over all images (frames) recorded in one sequence. To achieve a comparable standard for the calculation of the relative fluorescence changes we first calculated the ΔF/F_0_ and defined the background fluorescence (F_0_) as the mean of 10 successive frames at the beginning of each recording (frame 5 - 15). Second, we calculated relative values for ΔF/F_0_ in percent (%) which are plotted in the figures. For isolated brain imaging experiments each cluster response was normalized to its own strongest response, meaning that each cluster had a 100% value. Only fluorescence values of cluster crossing the half maximum threshold (50%) were taken into account for the wave pattern analysis. For *in vivo* imaging experiments cluster responses were normalized within each animal to its strongest response (100%) over all identified cluster, meaning that only one cluster had a 100% value. Normalized responses of VUM neuron cell body clusters were compared between test groups using unpaired t-tests with Welch’s correction for unequal variance when needed using statistical software (Prism, GraphPad Software, La Jolla, California, USA).

### Behavioral experiments

The foraging behavior of 3^rd^ instar larvae was investigated in glass dishes (13.5 cm Ø) which were cast using standard agarose (Axon Labortechnik, Kaiserslautern, Germany) in bidistilled water (2 mg/100 ml). The crawling events during normal foraging behavior and stimulus induced escape response of the animals were counted and their frequencies analyzed. Observations and mechanical stimulations were done at 26-27°C. For brush or rod stimulations either a handhold brush or a handhold rod stimulator (triggered by hand at the pulse generator) was used to stimulate the posterior end of the larvae. Quiescent animals or animals showing freezing or rolling behavior after mechanical stimulations were excluded from the analysis. Each experimental run started with the observation of the unstimulated foraging behavior during 20 seconds followed by the first stimulation. In total three successive stimulations with an ISI of 20 seconds were applied at 20, 40 and 60 seconds into the experimental run. After each stimulation the crawling events during 20 seconds were counted. For comparisons between different experimental groups a crawling index was calculated by dividing the number of crawling events after each stimulation through the number of crawling events of unstimulated animals before the first stimulation,

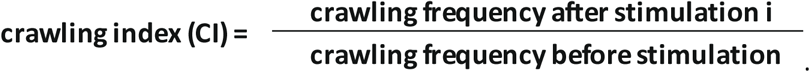

For correlation analysis between calcium imaging measurements and the behavioral performances we did linear regression analysis and calculated Pearson’s correlation coefficients (Fig. 4B).

### Immunohistochemical staining and confocal imaging

To visualize peripheral VUM neuron projections and innervation of body wall muscles in relation to neuronal muscular junctions (Fig. 4A) 3^rd^ instar larvae of Tdc2-GAL4/UAS-GCaMP3.0 flies were pinned on Sylgard dishes as described above. All organs were removed, and the body wall was fixed at six points using Minutien pins. The tissue was fixed for 10 minutes in 4% PFA (diluted in 0.1 M PBS), rinsed and washed several times in 0.1 M PBS-0.3% Triton X (PBS-Tx) and blocked 30 minutes in 10% normal goat serum (NGS, diluted in PBS-Tx). The blocking solution was exchanged with antiserum solution: rabbit-anti-GFP (Abcam 290, 1:1000) and mouse-anti-highwire (DSHB #6H4, 1:100) in 10% NGS. Tissue was incubated overnight at 4°C, washed 3 × 10 minutes in PBS-Tx and incubated for 2 hours (at room temperature) in second antiserum solution: goat-anti-rabbit-Alexa 488 (Invitrogen A-11008), goat-anti-mouse-Alexa 647 (Invitrogen A-21236) and Phalloidin-Alexa 594 (Invitrogen A12381). Tissue was washed in PBS and mounted in Vectashield (H-1000, Burlingame, Ca). To analyze central thoracic and abdominal VUM clusters (Suppl. Fig. 2A) larval brains (Tdc2-GAL4/UAS-GCaMP3.0) were dissected, fixed, washed and mounted in vectashield. Samples were stored at 4°C until being imaged with a Leica DMi8 CEL SP8 confocal microscope, 8-bit, 0.3 mm voxel depth (1.024 × 1.024 px). A 63x/1.40 HC PL APO Oil CS2 WD 0.14 mm objective was used. Images were processed with the freeware imaging software FIJI (Schindelin *et al.*, 2012) and the Leica Application Suite (LAS) X 3D analysis software.

## Author contributions

H.-J.P. and M.S. planned and designed the experiments. M.S., F.B. and M.-M.G. conducted functional calcium imaging experiments. A.S. performed confocal imaging. M.S., F.B. and M.-M.G. analyzed the data. H.-J.P. and M.S. drafted the manuscript.

## Acknowledgments

We express our gratitude to Heike Wolfenberg for excellent technical assistance and we thank Prof. Dr. Björn Brembs and Dr. Christine Damrau for providing fly lines. We thank Prof. Dr. Robin Hiesinger and Prof. Dr. Mathias Wernet for valuable discussion on the project. This work was supported by the German Federal Ministry of Education and Research (BMBF) through grant 01GQ1001D to the Bernstein Center for Computational Neuroscience Berlin.

