## Supplemental Captions for "Intensity coded octopaminergic modulation of aversive crawling behavior in *Drosophila melanogaster* larvae"

***Supplemental Figure 1. Animal preparation and imaging setup.*** **A** The CNS of 3rd instar Drosophila larvae were studied using calcium imaging techniques investigating VUM neuron activity in response to tactile stimulation. To gain access to the CNS the cuticle was dorsally cut in the first anterior third of the body and the gut, glands and fat body tissues were removed. A micro platform was used to stabilize the VNC. **B** Imaging setup and stimulation devises. A Polychrome II light source provided excitation light at 475 nm which was mirrored onto the preparation. The emitted light from GCaMP3 positive VUM neurons was captured by a CCD camera. Both, camera and light source were computer controlled and synchronized via the imaging control unit (ICU). Additionally, the two tactile stimulus devices (rod and brush stimulators) could be trigger by the imaging software. For gentle rod stimulations a loudspeaker (Ekulit 1517) membrane was deflected using a Grass (SD9) stimulator. For harsh brush stimulations the angle and movement of a servo motor holding the brush was controlled by an Arduino UNO (microprocessor developer board). Stationary movements of pinned larvae were video recorded by a CMOS camera from beneath the transparent experimental stage. **C** In the left image the experimental stage equipped with three micro manipulators can be seen holding the micro platform (lower left corner), the brush stimulator (upper left corner) and the rod stimulator (lower right corner). The opening in the center was holding the Sylgard dishes with the dissected animals. To the right in the same image you can see the Arduino connected to the brush stimulator. In the right image the experimental stage was installed between the microscope and a video camera capturing larval crawling movements.

***Supplemental Figure 2. Independent activity of individual VUM neurons.*** **A** reconstruction of abdominal VUM neuron cluster (a5-a9) based on confocal imaging sections. False color scaling represents scanning depth. Black arrow heads indicate individual cell bodies of VUM neuron cluster a7 and the white arrow head signify the location where the primary neurites bifurcate projecting laterally to both sides of the VNC. **B** In the upper and lower panel a series of 4 images (after 8, 14, 26 & 36 seconds and 5, 26, 44 & 48 seconds, respectively) with a close up view (40x objective) centered on abdominal VUM neuron cluster a1 and a2 (white circles) is shown. After pilocarpine superfusion of the preparation (5µl/1ml ringer; 10 minutes incubation) spontaneous fluorescence activity in individual cell bodies was measured. Cluster a1 and a2 show changes in fluorescence intensity of individual cell bodies a, b (blue and red arrow heads) and a, b and c (green, violet and light blue arrow heads), respectively. Line graphs in the middle represent changes in fluorescence intensity over time (in seconds) of individual cell bodies.

***Supplemental Figure 3. Behavioral crawling frequency after multiple tactile stimulations.*** Panel **A** and **B** are showing crawling frequencies (Hz) before stimulations (control) and after a $1^{st}$, $2^{nd}$ and $3^{rd}$ gentle rod or harsh brush stimulation, respectively. Based on these data we calculated the performance index used in bar graphs of figure 4C.

***Supplemental Movie 1. Forward wave pattern.*** Shown is a time lapse movie (speed x16, 5x loop) of a spontaneously occurring forward wave pattern in VUM neuron cluster in the isolated brain of a *Drosophila melanogaster* $3^{rd}$ instar larva (Tdc2-GAL4/UAS-GCaMP3.0).

***Supplemental Movie 2. Backward wave pattern.*** Shown is a time lapse movie (speed x16, 5x loop) of a spontaneously occurring backward wave pattern in VUM neuron cluster in the isolated brain of a *Drosophila melanogaster* $3^{rd}$ instar larva (Tdc2-GAL4/UAS-GCaMP3.0).

***Supplemental Movie 3. Unstimulated crawling larva.*** Shown is a 2 min time lapse movie (speed x5) with unstimulated stationary crawling movements during *in vivo* calcium imaging registration. Movies were recorded under infrared light from beneath the transparent microscope stage using a CMOS camera with its IR filter removed.

***Supplemental Movie 4. Pilocarpine response.*** Shown is a time lapse movie (speed x10) of VUM neuron activity in the VNC (all thorasic and abdominal cluster) after pilocarpine super fusion (5 min. incubation) using a 20x objective.

***Supplemental Movie 5. Pilocarpine response.*** Shown is a time lapse movie (speed x10) of VUM neuron activity in the VNC (cluster t3, a1-a3) after pilocarpine super fusion (10 min. incubation) using a 40x objective.

***Supplemental Movie 6. Brush stimulated larva.*** Shown is a time lapse movie (speed x2) of a dissected *Drosophila* larva under the microscope objective captured from beneath the transparent experimental stage. A harsh brush stimulation is applied to the posterior end of the larva during imaging recording of the VUM neuron cluster in the VNC.

***Supplemental Movie 7. Rod stimulated larva.*** Shown is a time lapse movie (speed x2) of a dissected *Drosophila* larva under the microscope objective captured from beneath the transparent experimental stage. A gentle rod stimulation is applied to the anterior end of the larva during imaging recording of the VUM neuron cluster in the VNC. The inserted window shows a close-up view of the rod's tip and the larval body wall which is deflected by a minute impulse.

***Supplemental Movie 8. Responses after repeated harsh brush stimulation.*** The time lapse movie (speed x10) in the upper panel shows an example of activity in the thorasic and abdominal VUM neuron cluster (t1-t3, a1-a5) in the VNC after posterior harsh brush stimulations (red spot in the sketch of the larva). Changes in fluorescence were measured over 60 seconds and three stimulations were applied after 10, 30 and 50 seconds, respectively (green bars in the line graph). The lower panel signifies relative changes in fluorescence in the VUM neuron cluster over time. The moving yellow bar in the line graph is synchronized with the grey scale movie. The data show increasing fluorescence responses after repeated stimulations with a lack of baseline recovery within 20 seconds from the second stimulation on.

***Supplemental Movie 9. Responses after repeated gentle rod stimulation.*** The time lapse movie (speed x10) in the upper panel shows an example of activity in the thorasic and abdominal VUM neuron cluster (t1-t3, a1-a5) in the VNC after posterior gentle rod stimulations (red spot in the sketch of the larva). Changes in fluorescence were measured over 60 seconds and three stimulations were applied after 10, 30 and 50 seconds, respectively (green bars in the line graph). The lower panel signifies relative changes in fluorescence in the VUM neuron cluster over time. The moving yellow bar in the line graph is synchronized with the grey scale movie. The data show decreasing fluorescence responses after repeated stimulations.
