## Supplementary figures and images for "Intensity coded octopaminergic modulation of aversive crawling behavior in *Drosophila melanogaster* larvae"

### Supplemental Figure 1

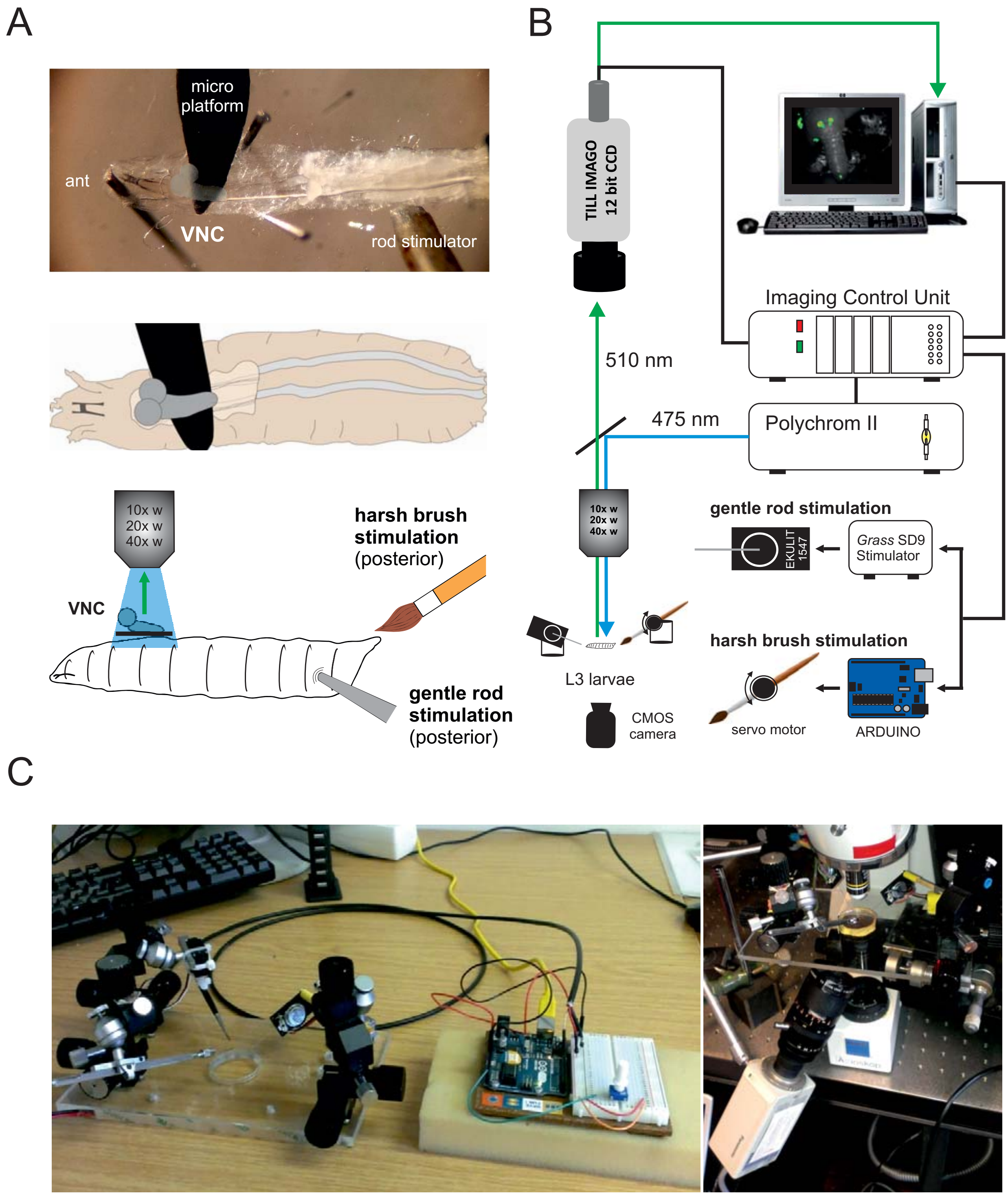

### Supplemental Figure 2

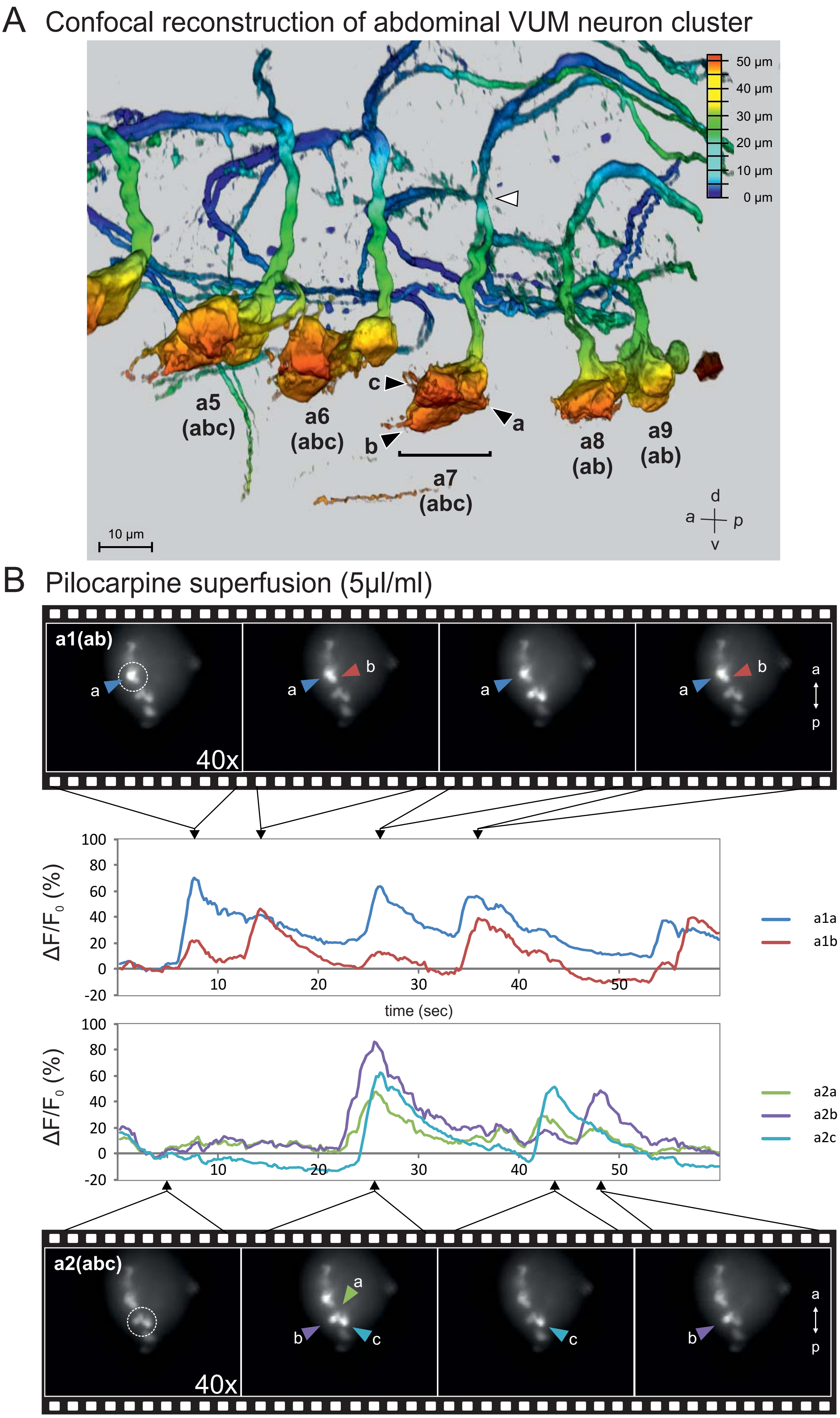

### Supplemental Figure 3

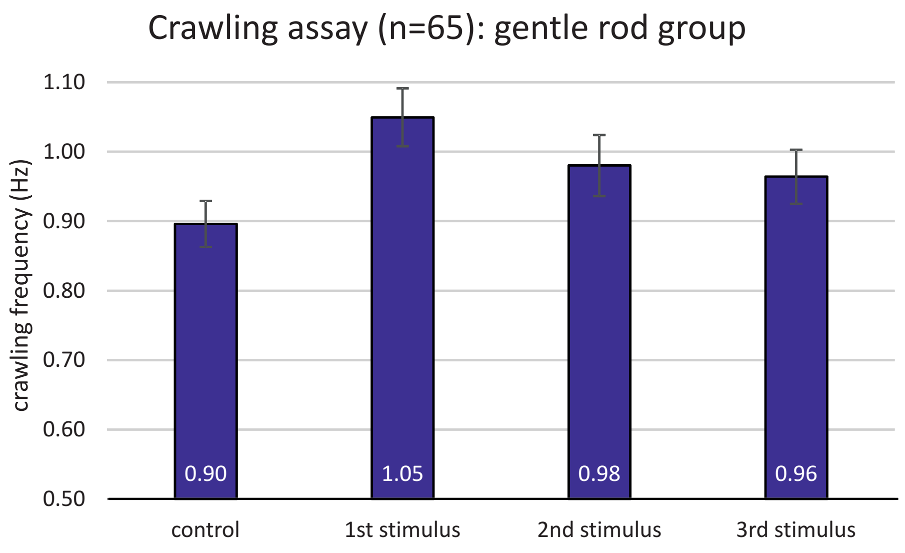

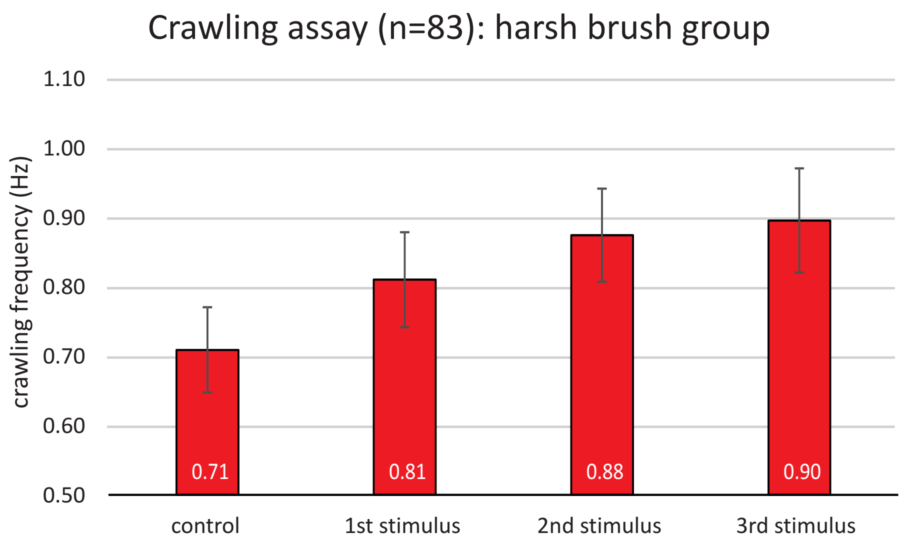
